# Optimizing Equilibration Time to Enhance Post-Thaw Viability of Cryopreserved Zebrafish (*Danio rerio*) Ovarian Fragments

**DOI:** 10.1101/2025.10.28.685221

**Authors:** Maya L. Wade, Matthew K. Litvak

## Abstract

Cryopreservation enables the long-term storage of viable biological material at ultra-low temperatures and forms the foundation for germplasm cryobanks that maintain valuable genetic lines of model organisms such as zebrafish (*Danio rerio*). However, reliable and reproducible cryopreservation protocols for fish germline stem cells remain difficult to develop, partly because key steps such as equilibration are often overlooked or assigned arbitrarily. Here, we optimized equilibration time for cryopreservation of zebrafish ovarian tissue. Ovarian fragments were equilibrated in 2 M methanol + 0.1 M glucose + 10% egg yolk for varying durations (15-120 minutes) before controlled slow-cooling and storage in liquid nitrogen. Post-thaw viability was assessed using a Trypan Blue exclusion assay. A 60-minute equilibration yielded the highest viability of ovarian cells in Experiment 1 (55.68 ± 1.74%), whereas a 30-minute equilibration yielded the highest viability in Experiment 2 (75.86 ± 2.44%), but was not significantly different from the 60-minute equilibration in Experiment 2 (75.58 ± 2.04%) (*p* = 0.9983). Equilibration alone accounted for a 48.71% increase in post-thaw viability relative to controls. The framework presented here provides a reproducible method for determining species-specific equilibration optima and supports the development of effective germplasm cryobanks for both model and endangered fish species.

## Introduction

Cryopreservation enables the long-term storage of viable biological material at ultra-low temperatures and serves as the foundational method behind germplasm cryobanks^1^. These cryobanks are essential for maintaining genetic lines of model organisms such as zebrafish (*Danio rerio*) and provide a framework for conserving the genetic diversity of endangered species^1^. At −196^°^C in liquid nitrogen (LN2), all biological activity stops, including the processes which cause cell death and DNA degradation^2,3^; thus, living material can be stored for extended periods without deteriorating (estimated 200 to 32,000 years for fish cells in LN2)^4^. When paired with assisted reproductive technologies such as germ cell xenotransplantation, germplasm cryobanks offer insurance against the loss of valuable strains or species while avoiding the logistical challenges of maintaining live captive populations^2,3,5–9^.

However, cryopreservation protocols that yield a high post-thaw viability of germ cells have proven difficult to develop. The most effective strategy for long-term preservation of genetic material would be cryopreservation of fish oocytes and embryos, but reliable and reproducible protocols have yet to be established despite recent experimental progress^1,3,10,11^. Sperm cryopreservation is the most established and commercialized technique, yet it only preserves the male germplasm, making it ill-suited for species that exhibit female heterogamy^12^. Cryopreservation of primordial germ cells (PGCs) would preserve both the maternal and paternal genomes and produce functional gametes via germline chimerism^8^. Nonetheless, endogenous PGCs are few in number (∼20 in zebrafish) and some are typically lost during isolation^13^. Germline stem cells, such as spermatogonia (SG) and oogonia (OG), offer a promising alternative because they are more abundant than PGCs but still retain considerable sexual plasticity and the ability to produce functional gametes via xenotransplantation^5,14,15^.

Post-thaw viability of cryopreserved germline stem cells varies widely across studies, reflecting both species-specific factors and differences in key protocol parameters^16^, such as the cooling regime, cryoprotectant formulation, equilibration time, extender medium, and thawing procedure. Table 1 summarizes previous zebrafish SG and OG cryopreservation studies, illustrating both the promise and inconsistency of current protocols. These findings highlight the need for systematic optimization of protocol steps to improve efficiency and effectiveness^12^.

**Table 1.**
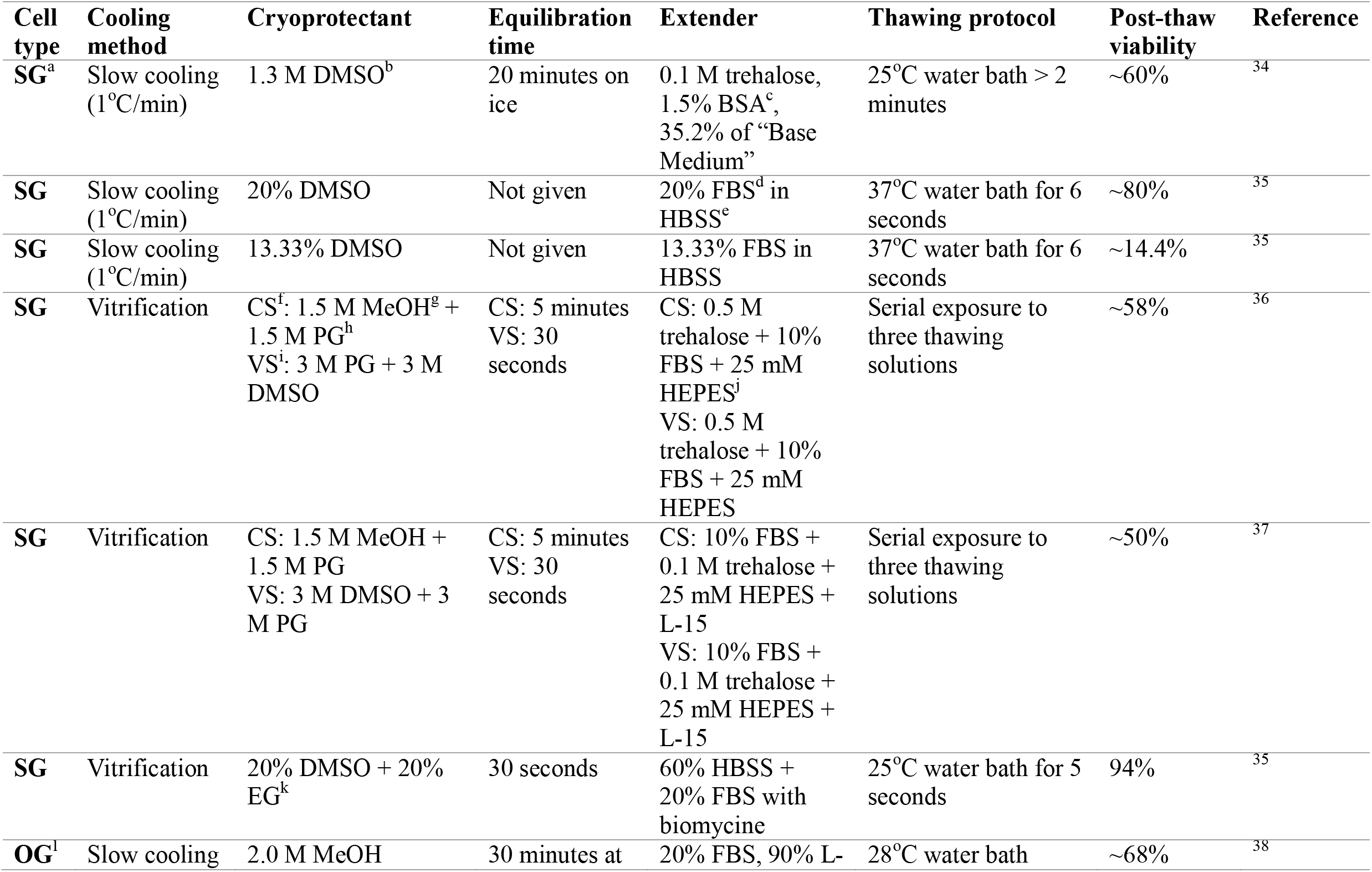
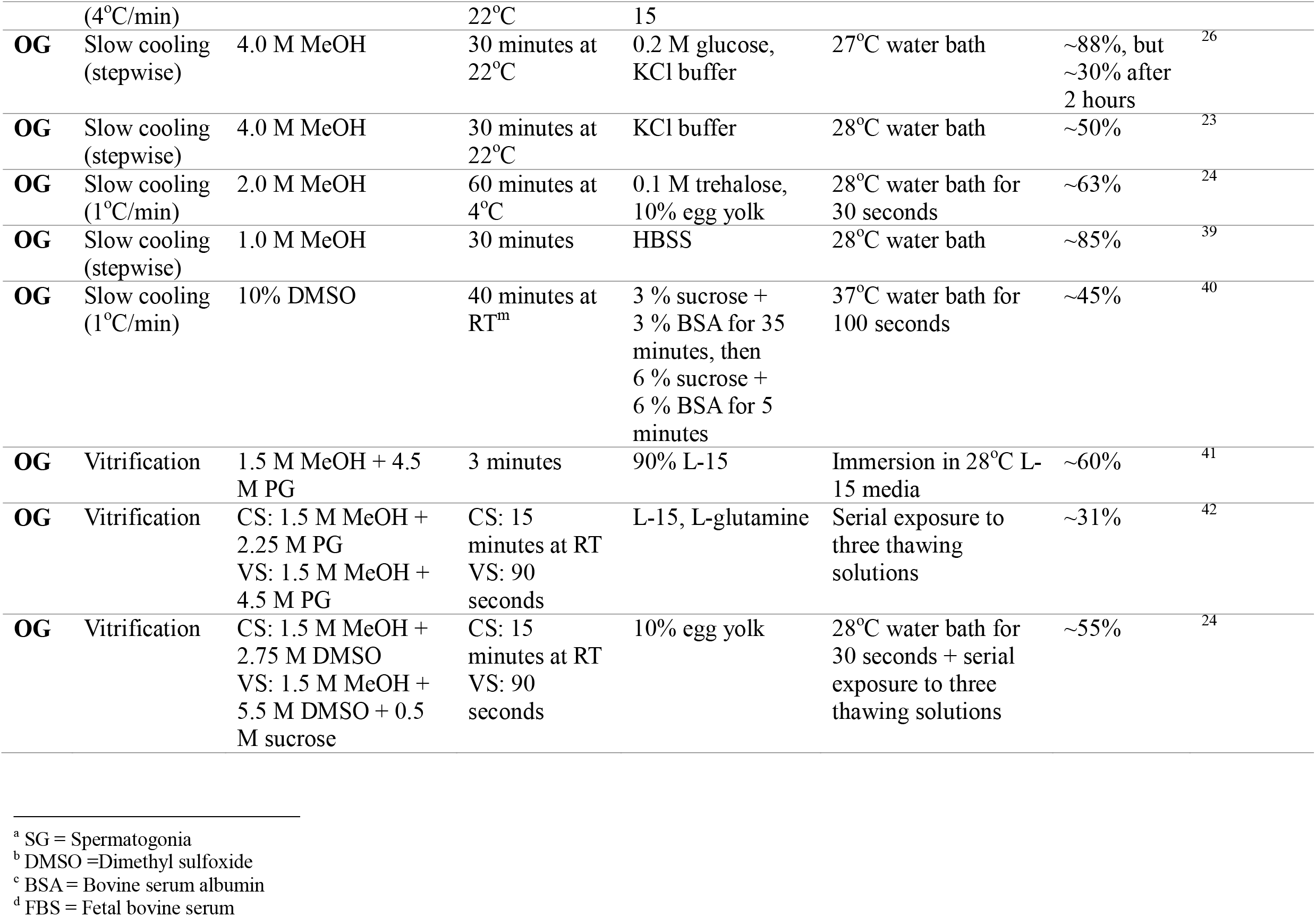
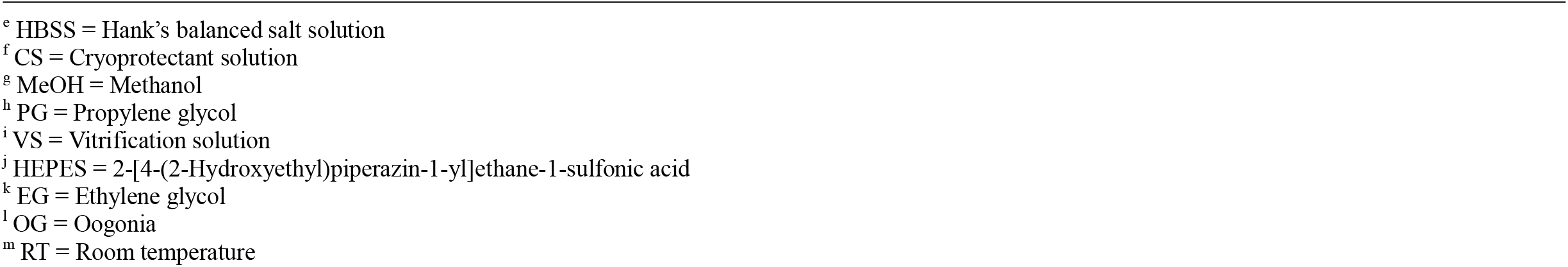
Review of zebrafish spermatogonial and oogonial stem cell cryopreservation studies, highlighting protocol differences and resulting post-thaw viability of germline stem cells.

The choice between cryopreserving SG or OG depends on the target species’ sex determination system. To retain both maternally and paternally inherited DNA, it is essential to preserve the heterogametic cell type. For example, in a female-heterogametic species such as the shortnose sturgeon (ZW/ZZ system), cryopreserving SG would result in the loss of maternally inherited DNA, which is not ideal for restorative efforts of an endangered species^17–19^.

The zebrafish, which also exhibits female heterogamy^20^, serves as a practical model for developing germline cryopreservation protocols for heterogametic fishes at risk. Its small size, rapid generation time, and high fecundity^21^ make it more tractable than fish like shortnose sturgeon, which is large-bodied, late-maturing, and endangered^22^. While previous studies have optimized cryoprotectant formulas^23–25^, cooling regimes^24^, and thawing methods^26^ for zebrafish ovarian fragments, equilibration time has yet to be optimized. Equilibration, the period during which the tissue is exposed to cryoprotectants before freezing, is a critical yet often overlooked step in the cryopreservation process^27^. Insufficient exposure to cryoprotectants leaves cells vulnerable to intracellular and extracellular ice formation, whereas prolonged exposure increases the risk of cryoprotectant toxicity and cell death even prior to freezing^16,28^. Therefore, optimizing equilibration time is essential for maximizing post-thaw viability and ensuring consistent cryopreservation outcomes.

In this study, we aimed to optimize the equilibration time for cryopreservation of zebrafish ovarian tissue. Specifically, we examine the effects of varying equilibration times on post-thaw cell viability so that the optimal equilibration time can be identified. These methods can be adapted to other fish species to determine optimal equilibration parameters during protocol development. Our findings contribute to the refinement of germplasm cryobanking practices for both model organism research and the conservation of endangered fish species.

## Materials and Methods

### Fish Husbandry

Zebrafish were purchased at PetSmart and maintained in a standalone recirculation multi-rack tank system (Aquabiotech Inc. Coaticook, Quebec, Canada). We used 2 racks equipped with 12 × 3 L tanks and 6 × 10 L tanks. Water was kept at 27°C, pH 6–7, and 0 ammonia. Water quality was monitored daily, with maintenance as needed. Fish were kept on a 14:10 hour light:dark cycle and fed three times daily with either freeze-dried tubifex worms (Omega One) or 1.0 mm slow-sinking pellets (Zeigler, adult zebrafish irradiated diet). Tanks were cleaned biweekly.) or 1.0 mm slow-sinking pellets (Zeigler, adult zebrafish irradiated diet). Tanks were cleaned biweekly.

### Ovarian Fragment Collection

Zebrafish were euthanized with a lethal dose of tricaine methane sulfonate (300 mg/L MS-222) buffered to neutral pH with sodium bicarbonate, followed by decapitation to ensure death. Total length was recorded prior to dissection. Ovaries were excised into 90% Leibovitz L-15 medium (Fisher Scientific) and cleared of fat under a light microscope. Ovaries were divided into five (experiment 1) or four (experiment 2) approximately equal-sized fragments using a scalpel and forceps, then kept on ice in 2.0 mL Eppendorf cryotubes containing 500 μL of 90% L-15 medium until cryoprotectant exposure.

### Cryoprotectant Solution Preparation

A cryoprotectant solution for zebrafish ovarian tissue was optimized by Marques *et al*.^24^ as 2 M methanol (MeOH) + 0.1 M trehalose + 10% egg yolk solution. Franěk *et al*.^29^ found no significant differences between supplementation with trehalose, fructose, or glucose, so we substituted 0.1 M trehalose for 0.1 M glucose (Fisher Scientific, 186122A). The 10% egg yolk was prepared as described by Marques *et al*.^24^. Fresh egg yolk was mixed with 90% L-15 medium (1:2 ratio) then centrifuged for 30 minutes at 10,000 rpm at 15^°^C in a temperature-controlled centrifuge (Sorvall, ST16R). Only the supernatant was added to the cryoprotectant solution, yielding a final composition of 2 M MeOH (VWR, BDH20864.400) + 0.1 M glucose + 10% egg yolk.

### Experiment 1 – Optimization of Equilibration Time

Ovarian fragments were transferred into 2.0 mL cryotubes containing 500 μL of cryoprotectant and gently mixed by pipetting. Within each individual, four tissue fragments were equilibrated on ice for 15, 30, 45, or 60 minutes. The fifth fragment served as a control and was held on ice for 60 minutes in 500 μL of 90% L-15 medium without cryoprotectant. After equilibration, samples were frozen in a PLANER Kryo-360 cryogenic freezer (cooling rate 1^°^C/min to -80^°^C) then stored in a LN2 dewar for at least two days.

Samples were thawed in a 28^°^C water bath (VWR, WB20) for 30 to 60 seconds (at a rate greater than 300^°^C/min) as optimized by Guan *et al*.^26^. A modified version of their cryoprotectant removal technique^26^ was used: samples were sequentially immersed in 1.0 M MeOH, 0.5 M MeOH, and 90% L-15 medium for 2.5 minutes each. Samples were gently pipetted to isolate cells, incubated in 0.2% Trypan Blue for three minutes, and rinsed three times in 90% L-15 medium to remove excess dye before visualization.

### Experiment 2 – Extended Equilibration Time Trial

We conducted Experiment 2 to determine whether equilibration longer than 60 minutes further improved post-thaw viability. Methods were identical to experiment 1 except that equilibration times were 30, 60, 90, and 120 minutes.

### Experiment 3 – Viability of Fresh Ovarian Tissue

Experiment 3 controlled for the freezing process. We used three zebrafish, each providing five ovarian tissue samples, to determine the viability of fresh (unmanipulated) ovarian cells. Freshly dissected fragments were stained with 0.2% Trypan Blue for three minutes and rinsed three times in 90% L-15 medium before visualization.

### Cell Visualization

We visualized cells using a Zeiss Axioskop 2 plus microscope and hemocytometer. Approximately 25 μL of stained sample was pipetted onto a slide and covered with a hemocytometer coverslip. Images were captured using the microscope’s imaging software to facilitate repeated counting. Cells were visualized under 2X and 10X objectives. A dual-tally counter was used to record stained (membrane-damaged) and unstained (membrane-intact) cells. Only samples with ≥ 100 cells were included in statistical analyses to avoid bias from low counts. Post-thaw viability was calculated as 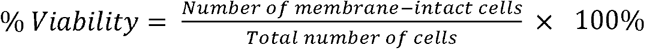.

### Statistical Analyses

We analyzed the effect of equilibration time (Experiments 1 and 2) on cell viability using repeated-measures ANOVA in R v4.0.0 “Arbor Day”. Code was adapted from Field *et al*.^30^ for a multilevel linear model. Normality and homogeneity of variances were assessed using Shapiro-Wilk and Bartlett tests, respectively, with significance set at α = 0.05. After verifying assumptions, we compared the multilevel linear model with a baseline model to evaluate whether inclusion of equilibration time as a predictor significantly improved model fit. Post-hoc comparisons were performed using Tukey contrasts for general linear hypotheses (*glht, multcomp* package^31^).

## Results

### Experimental Units

We used 41 female zebrafish (mean total length = 3.53 cm; 95% CI = 3.46–3.59 cm). In Experiment 1, samples from two zebrafish were lost due to cryoinjury, leaving undistinguishable cellular debris; these were excluded from statistical analyses. Final sample sizes were *n* = 27 (Experiment 1), *n* = 11 (Experiment 2), and *n* = 3 (Experiment 3).

### Experiment 1 – Optimization of Equilibration Time

Equilibration time significantly affected post-thaw viability (χ^2^(4) = 223.1, *p* < 0.0001). Post-hoc testing indicated that all groups differed significantly except between the 15- and 30-minute treatments (Figure 1). Viability increased with equilibration duration, reaching a maximum at 60 minutes 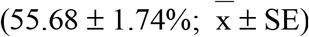. The average viability for the control was low (6.98 ± 1.60%).

**Figure 1.**
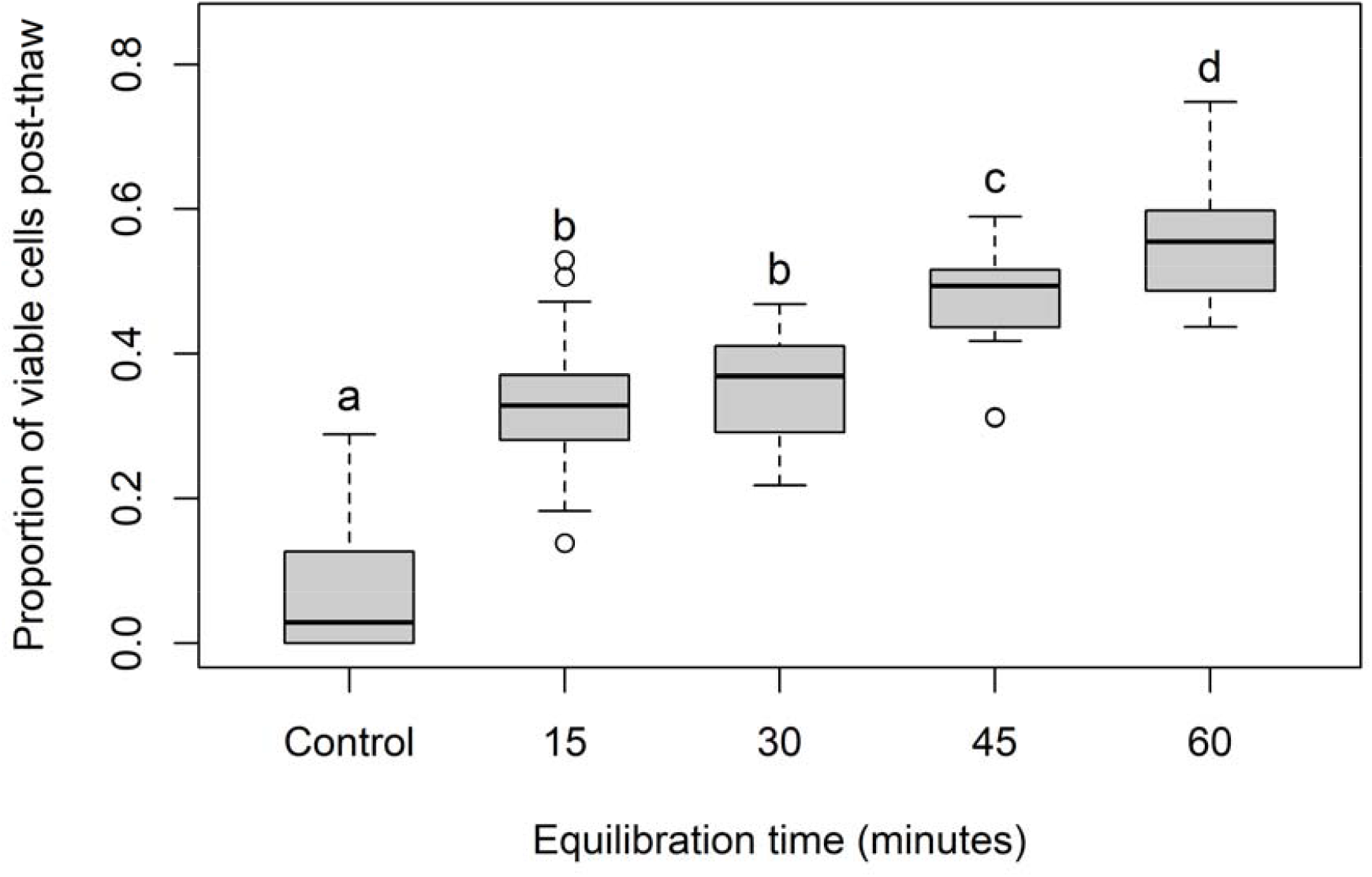
Post-thaw viability of zebrafish ovarian cells following different equilibration times (15, 30, 45, and 60 minutes). The control group was “equilibrated” without cryoprotectant. Equal letters denote no significant difference (*p* > 0.05).

### Experiment 2 – Extended Equilibration Time Trial

Equilibration time again significantly influenced post-thaw viability (χ^2^(3) = 20.4, *p* < 0.0001). The 120-minute treatment produced significantly lower viability than all other groups, whereas 30-, 60-, and 90-minute treatments did not differ significantly. As shown in Figure 2, the highest viability occurred between 30 and 60 minutes, although no solid conclusions can be drawn as differences between these groups were not statistically significant. Mean viabilities were 75.86 ± 2.44% (30 minutes), 75.58 ± 2.04% (60 minutes), 72.54 ± 2.24% (90 minutes), and 68.14 ± 1.95% (120 minutes).

**Figure 2.**
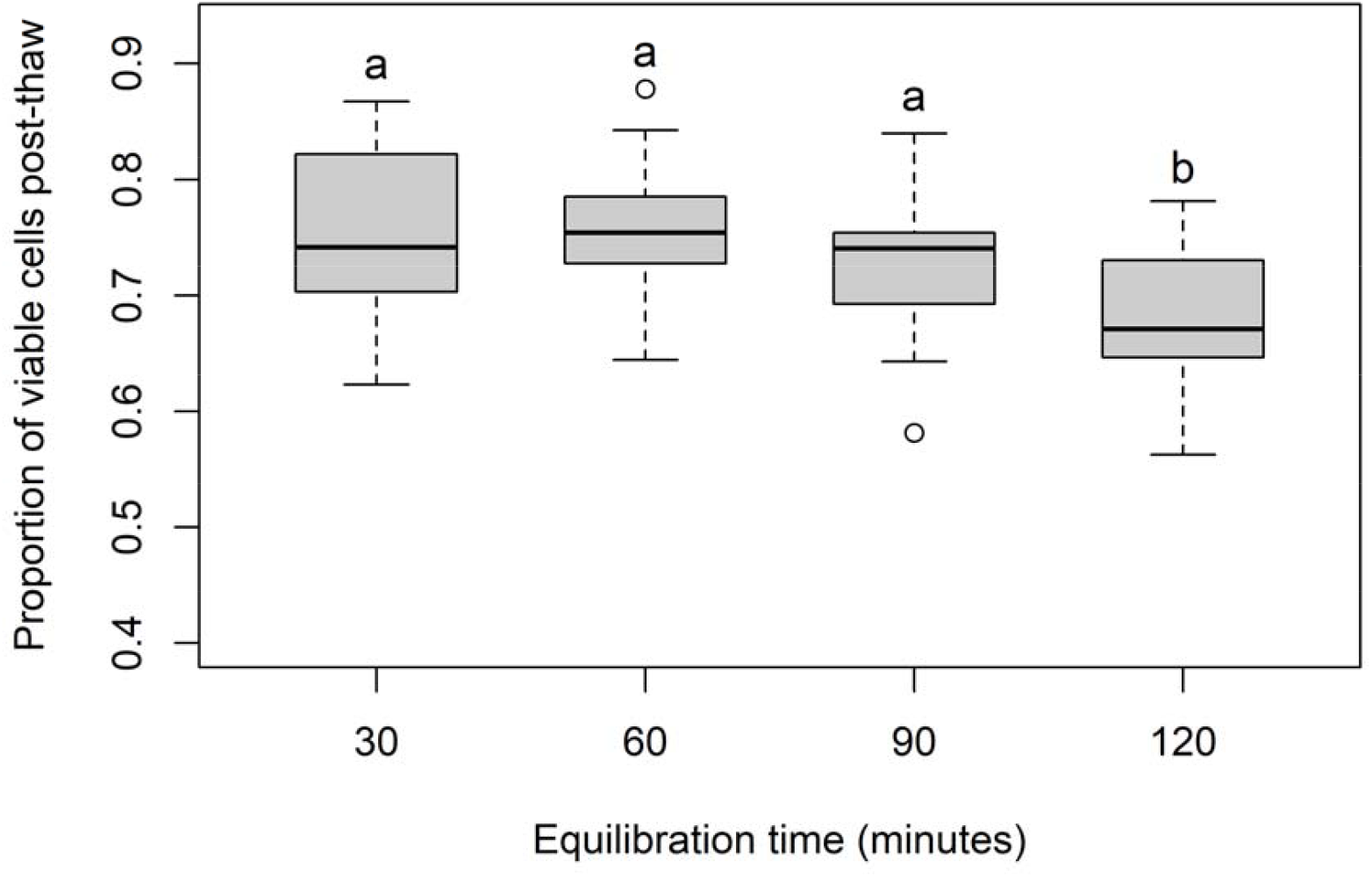
Post-thaw viability of zebrafish ovarian cells following extended equilibration times (30, 60, 90, and 120 minutes). Equal letters denote no significant difference (*p* > 0.05).

### Experiment 3 – Viability of Fresh Ovarian Tissue

Fresh ovarian cells exhibited high membrane integrity, with a mean viability of 95.98 ± 0.77%.

## Discussion

Cryopreservation of oocytes and embryos would provide a promising tool for the protection and propagation of endangered fish species. However, success has been limited by the large size, high lipid content, and polar organization of these cells^3^. For this reason, we focused on the cryopreservation of oogonial cells, which can differentiate into either female or male gametes when transplanted into the gonadal ridge of a sterilized fish. These germline stem cells are always present in fish gonads regardless of sex, age, or reproductive season^32^, making them reliable targets for germplasm preservation.

Our findings demonstrate that equilibration time is a critical determinant of post-thaw viability in zebrafish ovarian cells. In Experiment 1, viability increased with equilibration time, reaching a maximum at 60 minutes (55.68 ± 1.74%). The average viability for control samples was low (6.98 ± 1.60%), confirming the essential protective role of cryoprotectants. In Experiment 2, the highest mean viabilities occurred between 30 and 60 minutes (75.86 ± 2.44% and 75.58 ± 2.04%, respectively). Differences between these groups were not significant, likely reflecting limited statistical power (*n* = 11). Fresh ovarian tissue (Experiment 3) maintained high viability (95.98 ± 0.77%), indicating that dissection and handling contributed minimally (∼4%) to overall cell death.

Although logistical constraints during the COVID-19 pandemic prevented assessment of cell viability immediately after equilibration but prior to freezing, such data would have been valuable to isolate the effect of cryoprotectant toxicity from freezing injury. Future studies should include this intermediate step to determine which stage of the protocol results in the largest decrease in cell viability.

The lower viabilities observed in Experiment 1 compared to Experiment 2 (e.g., 60 minutes: 55.68 ± 1.74% vs. 75.58 ± 2.04%), likely reflect the smaller sample size of Experiment 2, inter-individual variation, and gradual improvements in our cryopreservation technique. Nonetheless, when data from both experiments were combined, a dome-shaped relationship between equilibration time and post-thaw viability emerged (Figure 3). Viability was low at short equilibration times, increased progressively to a maximum (the optimal equilibration time), and subsequently declined with longer exposure, likely due to cryoprotectant toxicity.

**Figure 3.**
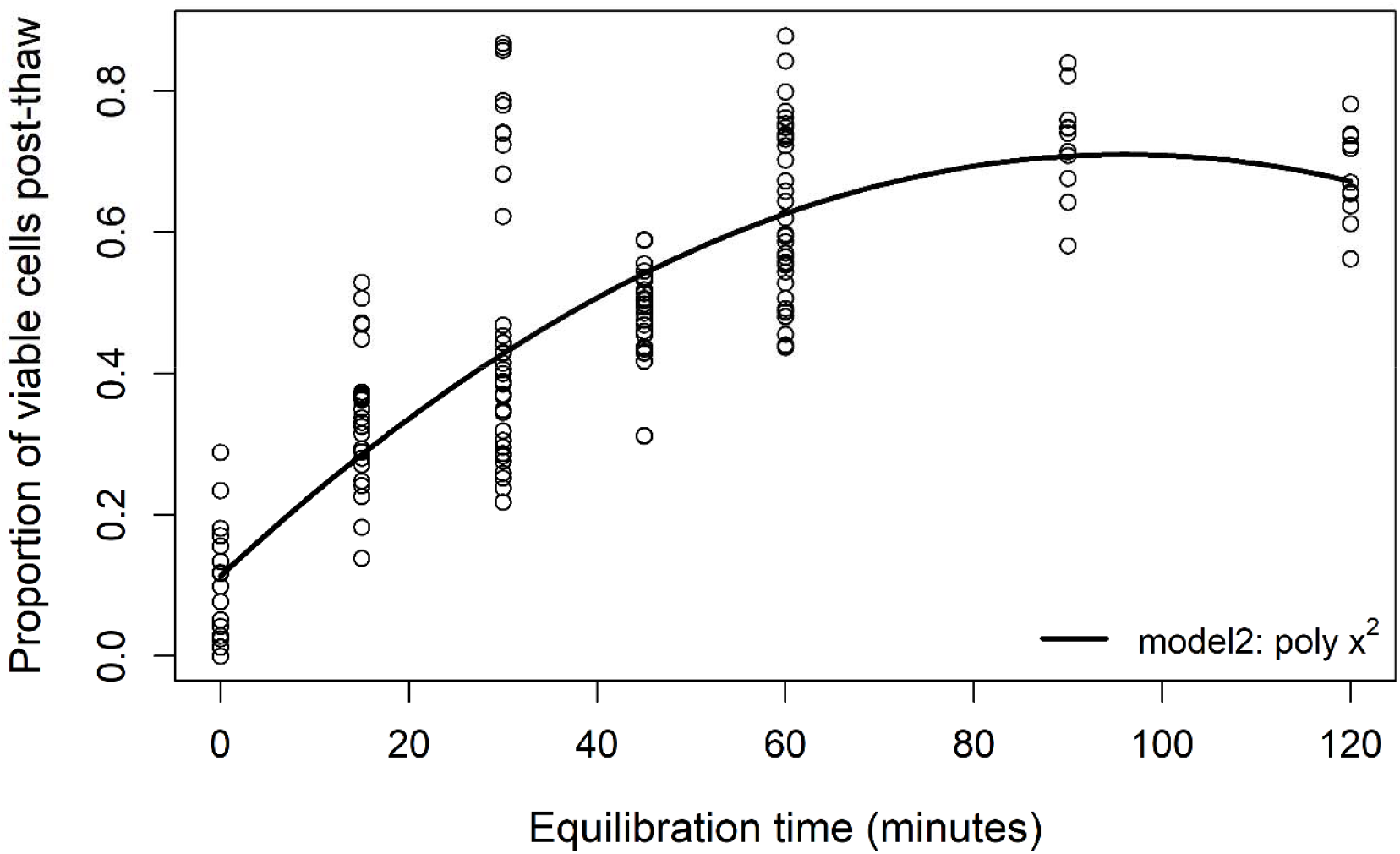
Combined data from Experiments 1 and 2 showing the effect of equilibration time on post-thaw viability of zebrafish ovarian cells. A polynomial regression (*R*^2^ = 0.6859) yielded the trendline (*y* = 0.1136 + 0.01244*x* – 0.00006484*x*^2^).

We performed a polynomial regression on the aggregated data to illustrate how an optimal equilibration time could be estimated. Using R version 4.0.0, a polynomial regression of the form *y* = *a* + *b*_1_*x* + *b*_2_*x*^2^ accounted for 68.59% of the variability in our response variable (*R*^2^ = 0.6859) and was tested against other models using an ANOVA. The resultant function with regression coefficients was *y* = 0.1136 + 0.01244*x* – 0.00006484*x*^2^. The optimal equilibration time in this example was 95.93 minutes with a viability of 71.03%. This analysis is illustrative only, as the aggregated dataset lacked a true repeated-measures design. With larger sample sizes and within-individual replication, a polynomial regression could be used predictively, and equilibration times near the modeled optimum could then be experimentally tested. Future work should therefore test a wider range of equilibration times within a single experimental design to develop a robust optimality response curve.

Our results are similar with those of Marques *et al*.^24^, who reported a maximum viability of ∼63.5% using a similar cryoprotectant solution (2 M MeOH + 0.1 M trehalose + 10% egg yolk) and a 60-minute equilibration period. In our study, the highest viability was 75.86 ± 2.44% using a 30-minute equilibration time (Experiment 2). However, we suspect the optimal equilibration time is closer to 60 minutes because if we combine the data from experiment 1 and experiment 2 the average viability of a 60-minute equilibration time is 61.60%, whereas the average viability of a 30-minute equilibration time is only 47.20%. More power from a larger sample size is needed to obtain a more accurate estimate of the optimal equilibration time.

To the best of our knowledge, this is the first study to explicitly optimize equilibration time for zebrafish ovarian tissue cryopreservation. Previous research has focused primarily on other protocol components^23–26,33^. Our findings underscore that equilibration is equally vital, as it governs how long cryoprotectants can interact with the tissue and provide protection from subsequent freezing. Despite its importance, equilibration time is often overlooked or chosen arbitrarily.

In our study, equilibration duration alone accounted for a 48.71% increase in post-thaw viability. Too little exposure to cryoprotectants leaves cells vulnerable to freezing injury, while excessive exposure risks cryoprotectant toxicity. Between these extremes lies an optimal “sweet spot” that maximizes post-thaw viability. Our results suggest that this optimum likely occurs near 60 minutes for zebrafish ovarian tissue, though additional refinement is needed. Given the importance of germplasm cryobanking for conservation, these findings provide a practical framework for developing species-specific equilibration protocols to improve cryopreservation outcomes for both model organisms and endangered species.

## Conclusions

This study demonstrates that equilibration time is a critical determinant of post-thaw viability in zebrafish ovarian tissue cryopreservation. Across experiments, viability increased with longer equilibration durations up to approximately 60 minutes, after which extended exposure to cryoprotectants reduced viability. These results highlight the importance of optimizing this often-overlooked step to improve overall cryopreservation outcomes.

Although further refinement is needed, particularly with larger sample sizes and a true repeated-measures design spanning a wider range of equilibration durations, the methods presented here provide a reproducible framework for determining species-specific equilibration optima. This approach can be readily adapted to other fish species to enhance the development of efficient, reliable cryopreservation protocols. By advancing the foundational methods required for germplasm cryobanking, this work contributes to broader efforts aimed at safeguarding aquatic biodiversity and supporting the propagation of threatened fish populations through assisted reproductive technologies.

## Acknowledgements

We would like to thank our funding agencies for their support of this study. We would also like to thank the MtA Litvak Lab research assistants and lab technicians who helped with this research, especially Seth Holditch, Lia Massoeurs, and Sophie Hartlen.

## Authorship Confirmation/Contribution Statement

**Maya L. Wade:** Methodology, Software, Validation, Formal Analysis, Investigation, Writing – Original Draft, Writing - Review & Editing, Visualization, Project Administration.

**Matthew K. Litvak:** Conceptualization, Methodology, Resources, Data Curation, Writing – Review & Editing, Supervision, Project Administration, Funding Acquisition.

